# Working memory availability affects neural indices of distractor processing during visual search

**DOI:** 10.1101/295378

**Authors:** Dion T. Henare, Jude Buckley, Paul M. Corballis

## Abstract

Working memory and selective attention are traditionally viewed as distinct processes in human cognition. However, increasing research demonstrates significant overlap between these constructs such that as working memory availability decreases, individuals perform worse on attention-based tasks. To date, the neural mechanisms involved in this interaction are unknown. We measured three candidate lateralized event-related potential components (N2pc, Ptc, and SPCN) to observe the effects of increased working memory load on selective processing of targets and distractors. We found that increased working memory load impaired the processing of distractors, but not targets, and this was reflected in attentuation of the Ptc to distractors. We also found that individual performance on the task is related to the neural response to both targets and distractors. This study suggests that working memory availability impacts individuals’ ability to disengage from irrelevant stimuli, and that individual differences in visual search ability under load are related to both target and distractor processing.

## Introduction

Everyday situations such as grocery shopping and driving depend on our ability to both efficiently allocate attention to relevant objects while suppressing distraction, as well as maintain and retrieve information from working memory. Although research over the past decade has demonstrated significant functional overlap between attention and working memory, the multifaceted nature of the relationship between these two constructs is not well understood (De Fockert, 2013; Fougnie, 2008). Converging evidence suggests that when load on a concurrent working memory task is increased, performance on an attention task is subject to increased interference from irrelevant distractors (Ahmed & De Fockert, 2012; Fockert, Rees, Frith, & Lavie, 2001; Stins, Vosse, Boomsma, & De Geus, 2004). There is also evidence that individuals low in working memory capacity show greater interference on a standard Stroop task (Kane & Engle, 2003), as well as greater interference on an Eriksen flanker task (Shipstead, Harrison, & Engle, 2012). Taken together, these findings suggest that the relationship between selective attention and working memory may be better framed in terms of working memory availability. From this perspective, increasing the working memory load on an individual would lead to a decrease in their working memory availability. What’s more, individuals with low working memory capacity would have less overall available working memory (see De Fockert (2013) for a review).

Contemporary accounts of working memory availability now typically incorporate attentional control either as an important contributor to working memory capacity(Kane, Bleckley, Conway, & Engle, 2001; Shipstead, Lindsey, Marshall, & Engle, 2014), or in some cases as the primary determinant of working memory capacity (Engle, 2018). The question of why working memory availability is important for selective attention remains unanswered. The aim of this research was to specify the relationship between selective attention and working memory by recording electrophysiological indices of attentional control under varying working memory load.

Optimal performance of selective attention relies on several underlying processes including; target identification and selection, proactive suppression of possible distractors, and e˚cient disengagement from irrelevant distractors that capture attention (Carrasco, 2011). Evidence for the effects of low working memory availability on attentional interference is equivocal, and interference with any one of these processes would produce impaired behavioral performance associated with low working memory availability. Some evidence suggests that low working memory availability delays the speed with which individuals can deploy attention to relevant targets (Heitz & Engle, 2007; Scalf, Dux, & Marois, 2011). Yet, findings from other work suggests that low working memory availability may cause an increase in capture by irrelevant distractors (Fukuda & Vogel, 2011; Vogel, McCollough, & Machizawa, 2005).

A different account was proposed by Fukuda and Vogel (2011) in research using both psychophysical and electrophysiological measures of recovery from attentional capture in low and high capacity individuals. Findings from this work suggest that the two groups are equally prone to initial capture of attention, but availability of working memory allows more efficient disengagement from irrelevant objects after they have captured attention.

The processes of target selection, distractor suppression, and distractor disengagement are difficult to tease apart using only behavioral measures. Nonetheless, recent research has identified a set of lateralized event-related potential (ERP) components that may provide dissociable measures of these processes. The first of these is the N2pc component, defined as an increased negative voltage at posterior electrodes contralateral to an attended stimulus. Accumulating evidence suggests that the N2pc indexes the shift of visual attention onto an attended object (Dell’Acqua, Sessa, Jolicoeur, & Robitaille, 2006; Hickey, McDonald, & Theeuwes, 2006; Luck & Hillyard, 1994). N2pc amplitude has also been used as a measure of attentional capture by relevant stimuli (Kiss, Jolicoeur, Dell’Acqua, & Eimer, 2008), or pop-out objects (Eimer & Kiss, 2007; Holgian, Doallo, Vizoso, & Cadaveira, 2009). In the present study, we took advantage of the contralateral control method in order to measure the N2pc elicited separately by targets and distractors (Hickey, Di Lollo, & McDonald, 2009; Hilimire & Corballis, 2014; Hilimire, Mounts, Parks, & Corballis, 2011). This method provided us with independent measures of N2pc to targets, reflecting target selection, and N2pc to distractors, reflecting initial distractor capture.

The second identified lateralized ERP component is an enhanced positivity contralateral to distracting or irrelevant objects (Hickey et al., 2009; Hilimire, Mounts, Parks, & Corballis, 2009; Toffanin, Jong, & Johnson, 2011). Research by Hickey et al. (2009) concluded that this component was a positivity related to distractor processing, thus they referred to it as “Pd”. Findings from research by Hilimire, Hickey, and Corballis (2012) suggest that the component reflects distractor suppression during the disambiguation of target features. A similar (or possibly identical) positive component referred to as “Ptc”, to reflect its temporal and contralateral scalp distribution has been identified in research by Hilimire et al. (2009). Findings from research by Hilimire et al. (2011) suggests that Ptc, like Pd, appears to reflect distractor suppression. However, evidence that both Pd and Ptc can be elicited by target objects suggests that Pd and Ptc may more generally reflect attentional disengagement from an attended stimulus (Burra & Kerzel, 2014; Hilimire & Corballis, 2014; Sawaki, Geng, & Luck, 2012).

In this study we measured lateralized ERPs separately for targets and distractors during the delay period of a working memory task. By varying the load on the working memory portion of the task, we were able to test specific predictions regarding the attentional mechanism that is affected by reduced working memory availability. Given that N2pc reflects attentional processing of an object, the effect of working memory availability on target selection was predicted to be reflected in modulations of the target N2pc. The effect of working memory availability on distractor capture on the other hand was predicted to be reflected in modulations of the distractor N2pc. Finally, given that Ptc reflects stimulus disengagement, the effect of working memory availability on distractor disengagement was predicted to be reflected in modulation of the distractor Ptc under varying working memory load.

## Methods

### Participants

22 individuals (8 males) between the ages of 18 and 29 (M = 22.09 years, SD = 3.13) participated in the experiment and were reimbursed with a NZD $10 voucher. All participants reported that they had normal or corrected-to-normal vision. All participants gave written, informed consent prior to the start of the experiment and the research was approved by the University of Auckland Human Participants Ethics Committee.

### Procedure

Testing took place in a dimly lit, electrically shielded chamber. Experimental stimuli were presented on an LCD computer monitor (Samsung Sync Master P2270), with screen dimensions of 47.7 × 26.8cm and a resolution of 1920 × 1080 pixels. The participant was positioned 57cm away from the monitor with the viewing distance maintained through use of a chinrest. E-prime software (version 2.0.8.74, Psychology Software Tools) was used to control stimulus presentation, synchronize stimulus presentation with the electrophysiological recording, and record participant responses. Participants performed a dual working memory task and a selective attention task in which they encoded either one or two shapes at the start of a trial that they had to match to a test display after some delay. During the delay period of the working memory task, participants performed 4 visual searches in a series of target-decoy displays. Each block consisted of 8 trials (8 working memory trials, 32 visual searches) and all participants completed a practice block followed by 18 testing blocks. This resulted in a total of 144 working memory trials and 576 visual search trials for analysis. Participants had a minimum 30 second break between each block and on half of the breaks they could choose to take a longer break.

Each trial began with a fixation cross on screen for 1500ms, and then the working memory cue appeared for 1000ms. In half of the trials, the cue would point either up or down (with equal probability) resulting in a low load trial, and in the other half of trials the cue would point both up and down resulting in a high load trial. The working memory encode display would then be presented for 2000ms and participants had to encode the shape(s) indicated by the cue.

At this point, the visual search task would begin, first with a fixation cross presented for between 500 and 1500ms (randomly selected on each trial), followed by the presentation of the visual search display which appeared for 200ms. Participants had 1600ms to respond to the orientation of the target letter. If the colored T was upright they would press “1” on a standard keyboard numpad with their right hand, whereas if the T was rotated 180^°^ then they would press “2”. Participants were instructed to respond as quickly as possible while attempting to maintain their accuracy above 90%. The visual search portion of the task was repeated 4 times in each trial.

Once the 4th visual search was completed, the working memory retrieval display would appear on screen for 2000ms and participants had to press “m” with their left hand if the shape(s) on screen matched the encoded shape(s), or “x” if it did not. The shapes matched on half of the trials. Finally, participants received feedback on the working memory portion of the task with a 1000ms display of either “Correct!”, “Wrong”, or “Too Slow” depending on their response. For an example trial, see figure 1.

**Figure 1.**
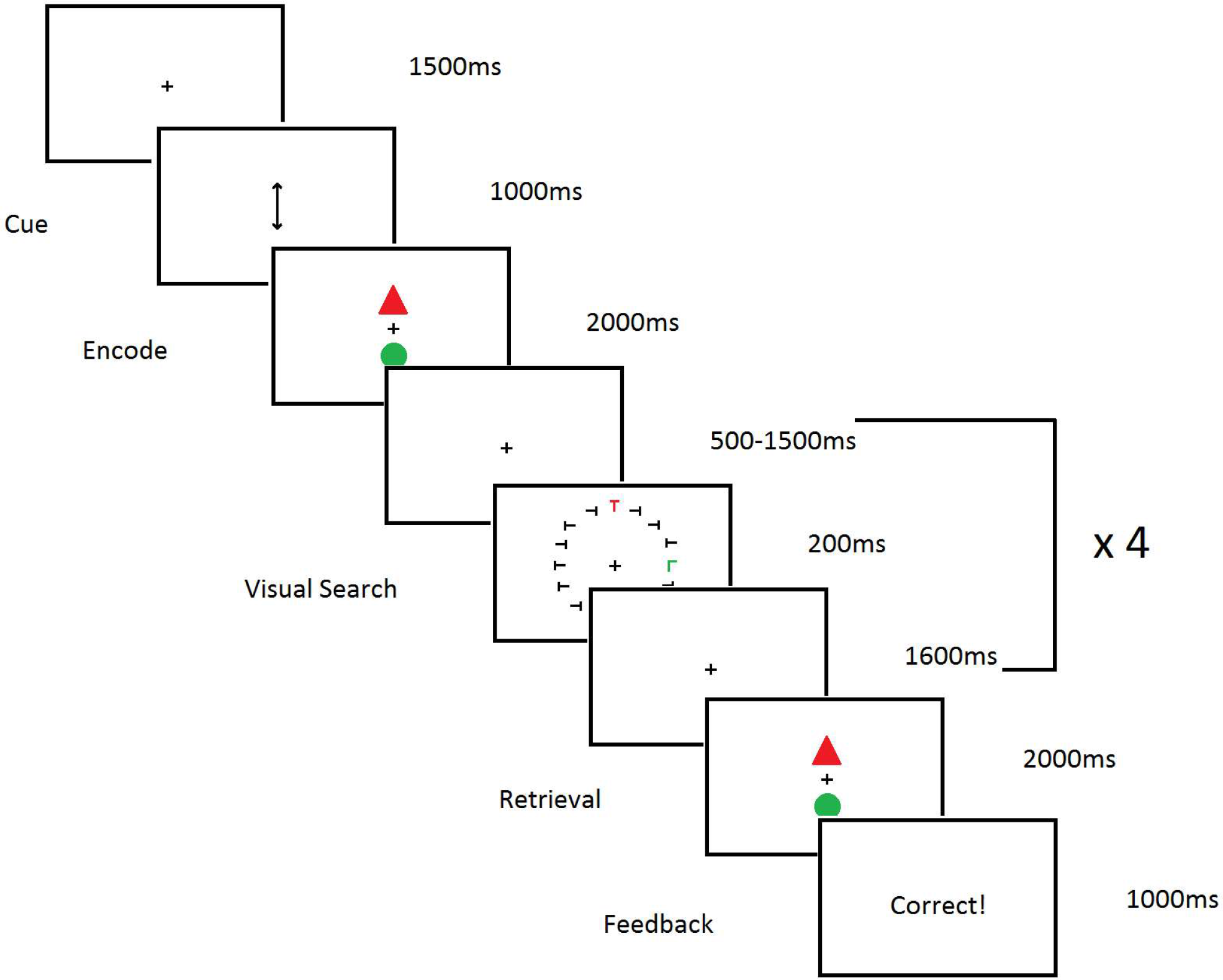
Example of a single trial of the experiment. This example shows a high load condition, as indicated by the double headed arrow cue pointing to both locations. In a low load trial the arrow cue would point at only one of the possible locations.

### Stimuli

All stimuli were presented on a black background, and a gray fixation cross (0.2^°^ x 0.2^°^) was maintained in the center of the screen throughout the trial. The working memory cue consisted of a gray vertical arrow (0.4^°^ x 1.5^°^) which replaced the fixation cross and pointed either up, down, or both up and down. The working memory encoding and retrieval displays consisted of two colored shapes (each 1.2^°^ x 1.2^°^) presented on the midline, one centered 2.5^°^ above and one centered 2.5^°^ below the fixation cross. These stimuli could be one of four shapes (triangle, square, circle, diamond) in one of four colors (green, blue, orange, pink) however the two stimuli were never the same shape or color on a given trial. The visual search displays consisted of 16 letters (Ts and Ls; 0.6^°^ x 0.6^°^) arranged in equal intervals around an imaginary circle centered on the fixation cross with a radius of 3^°^. Fourteen of the letters were gray Ts randomly rotated 90^°^ to either the left or the right. The remaining two letters were a colored T (the target) and a colored L (the distractor). The target was randomly selected to be either green or orange on each trial, and the distractor was always the color not selected for the target. The target and distractor were randomly selected to be either upright or rotated by 180^°^ on each trial, independent of one another. One of these two objects was always placed on the midline (half of the time the target, half of the time the distractor) while the other was lateralized to the left or right but always in the same visual field (upper or lower) as the midline object. This ensured that there was only ever one salient lateralized object for the calculation of lateralized ERPs.

### Electrophysiological recording

EEG activity was recorded continuously at 1000Hz on an Electrical Geodesics Inc. (EGI) Hydrocel-GSN-128-channel Ag/AgCl electrode net (Electrical Geodesics Inc.,Eugene, OR, USA). The electrodes have a geodesic arrangement and extend inferior to the axial plane containing F9/10 in the 10-10 system. The online reference electrode is Cz (placed on the vertex) and a COM sensor is located on the z axis between Pz and POz equivalents. EEG activity was amplified with an EGI net amps 400 amplifier with 24-bit A/D conversion, an input impedance of ≥ 200 MΩ, and a 4kHz antialiasing filter (500Hz low pass cut off). Electrode impedance levels were kept below 40kΩ and EGI’s implementation of the driven common technique (Common Active Noise Sensing and Limiting) was used throughout.

### EEG Processing

EEG data were processed twice, firstly in a way which was optimized for the performance of independent component analysis (ICA), and secondly in a way which was optimized for the production of ERP components. ICA weights from the first set of data could then be copied across in order to perform artifact correction in the ERP data. All EEG preprocessing was performed using MATLAB (2017a, The Mathworks Inc.) and the EEGLAB toolbox (Delorme & Makeig, 2004).

For the performance of ICA, data were down-sampled to 250 Hz. Data were Band pass filtered from 1 to 30Hz using a butterworth filter implemented in ERPLAB (Lopez-Calderon & Luck, 2014). Channel locations were loaded. Bad channels were detected, removed, and replaced with a spherical interpolation of surrounding electrodes. Data were then re-referenced to the average of all electrodes. ICA was then performed on the data, using the PCA option to reduce dimensions to account for the number of interpolated channels. ICA weights were then stored in order to be applied to data which had been pre-processed for the production of ERPs. For the production of ERPs, data were first down-sampled to 250Hz. Data were Band pass filtered from 0.1 to 30Hz using a butterworth filter implemented in ERPLAB (Lopez-Calderon & Luck, 2014). Channel locations were loaded. Bad channels were detected, removed, and replaced with a spherical interpolation of surrounding electrodes. Data were then re-referenced to the average of all electrodes. Data were then epoched from 200ms pre-stimulus to 800ms post-stimulus. Epochs containing horizontal eye movements were removed (identified as any epoch containing a greater than 32 microvolt difference between electrodes location at the outer canthi of each eye). ICA weights were then copied over to this set of data in order to perform artifact correction. The ADJUST toolbox was used in order to identify artifact components (Mognon, Jovicich, Bruzzone, & Buiatti, 2011). All components identified as artifacts by ADJUST were removed from the data. Subsequently, epochs which contained any activity above or below 100 microvolts were removed from the data. Participants with less than 60% of their trials remaining at this point were rejected, resulting in 2 rejections.

In order to isolate activity elicited contralateral to the lateralized object, all trials containing a right visual field target or distractor had their EEG activity flipped along the longitudinal axis such that for all trials, right hemisphere electrodes now represent activity contralateral to the lateralized stimulus and left hemisphere electrodes represent activity ipsilateral to the lateralized object. Ipsilateral activity was then subtracted from both hemispheres and the remaining contralateral activity was then mirrored onto the left hemisphere. At this stage, scalp activity represents only that portion of the EEG signal which is greater contralateral to the lateralized stimulus, mirrored onto both sides of the scalp. Importantly for our results, this means that in target lateralized trials (where the distractor is on the midline), activity elicited by the distractors is subtracted out of the data and only target related activity remains. Conversely, in the distractor lateralized trials (where the target is on the midline), activity elicited by targets has been removed.

Participant averages were formed separately for lateralized target and lateralized distractor trials and global field power (GFP) was then calculated for each of these conditions (across all electrodes posterior to Cz). In order to find unbiased estimates of the component latencies and their widths, Matlab’s built-in findpeaks function was used to detect the five most prominent peaks in the GFP for each stimulus type as well as their widths (using half-prominence). If any of these peaks overlapped with the pre-stimulus period then they were ignored, leaving 4 components of interest for both the target and the distractor lateralized conditions. Electrode clusters were then selected for each component by averaging across working memory load and plotting the topographies for each window. Electrode selection was similar for all components and centered around P7/8. For the targets, an early positivity (156 to 180ms), followed by three negativities (208 to 252ms, 380 to 488ms, and 676 to 700ms). The latencies and scalp distributions of these components line up with the P1pc, N2pc, and two parts of the SPCN component, as a result we will refer to them as the P1pc, N2pc, SPCNa, and SPCNb (shown in figure 3). For the distractors, this method identified an initial negativity (208 to 256ms), followed by a positivity (296 to 360ms), followed by two negativities (532 to 596ms, and 700 to 728ms). These latencies and scalp distributions are consistent with the N2pc, Ptc, and two portions of the SPCN component and we will therefore refer to them as the N2pc, Ptc, SPCNa, and SPCNb (shown in figure 2). For the individual diffrences analyses we used a signed-area approach similar to that used by Gaspar, Christie, Prime, Jolicoeur, and McDonald (2016) and Sawaki et al. (2012). First we calcuated the positive and negative area across all components for targets and distractors under high and low load. In order to correct for the effect of noise in these estimates we followed the procedure of Gaspar et al. (2016) and subtracted the equivalent calculation taken from the prestimulus area in each condition (and in our case this value was multiplied in order to account the different lengths of time between the prestimulus period and the area of the components).

**Figure 2.**
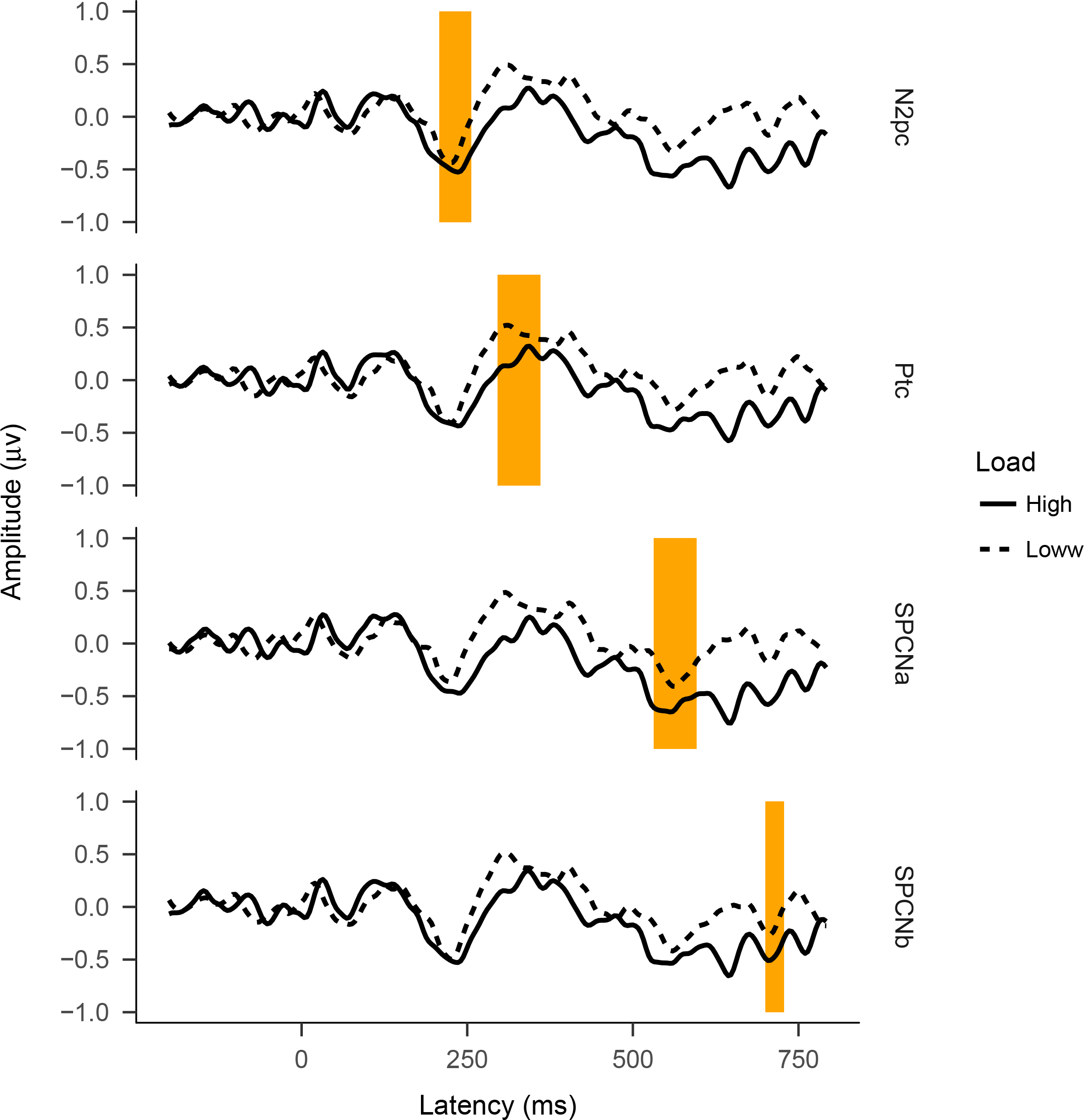
Subtracted event-related potentials showing the lateralized response to distractors under high and low load. Each component was calculated from the average of a slightly different cluster of electrodes (based on the condition averaged scalp topography of that component’s time window).

**Figure 3.**
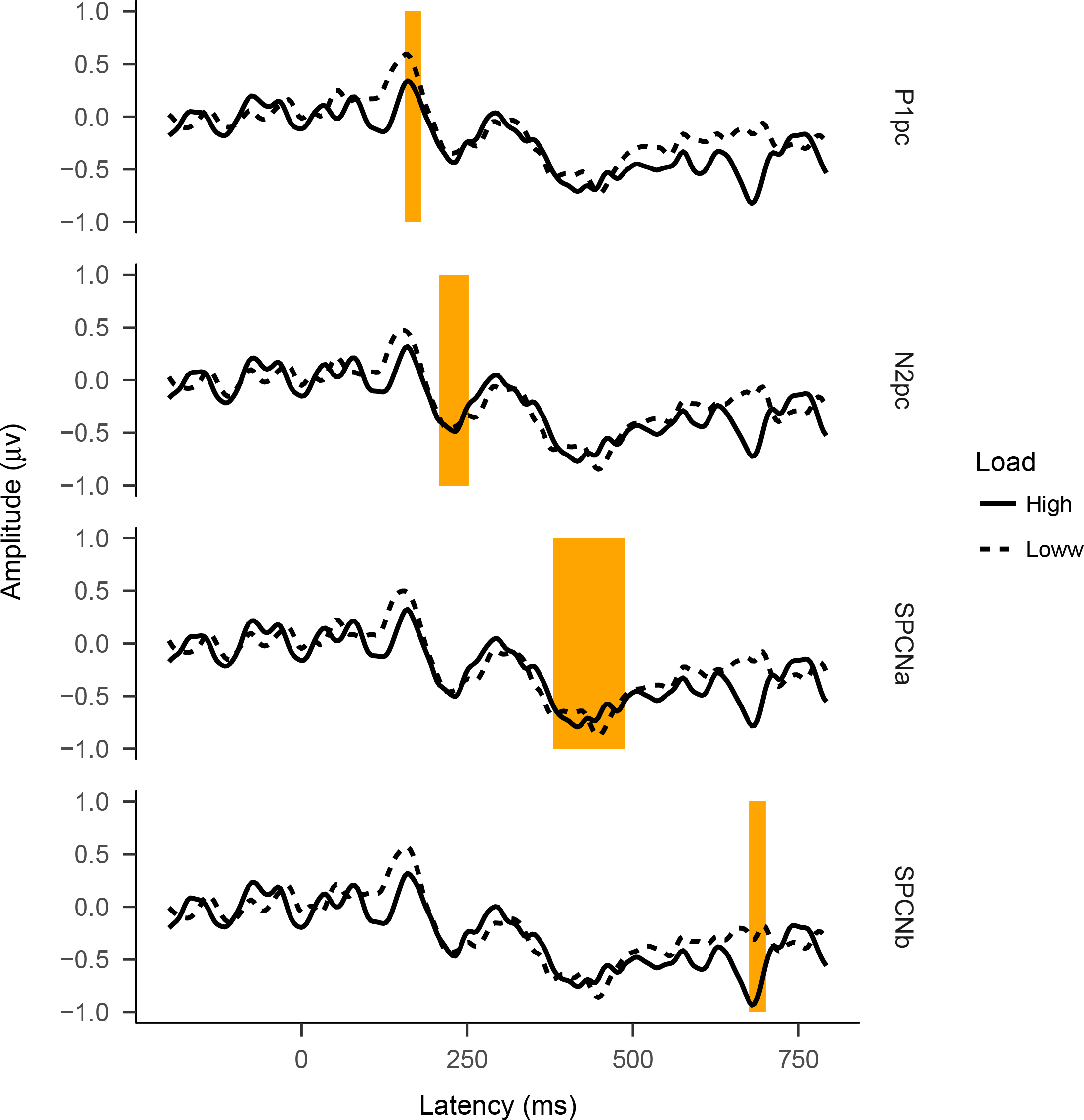
Subtracted event-related potentials showing the lateralized response to targets under high and low load. Each component was calculated from the average of a slightly different cluster of electrodes (based on the condition averaged scalp topography of that component’s time window).

### Statistical analysis

All statistical anaysis was performed in version 1.1.423 of RStudio (RStudio Team, 2016) using R (Version 3.4.3; R Core Team, 2017) and the R-packages *afex* (Version 0.19.1; Singmann, Bolker, Westfall, & Aust, 2018), *bindrcpp* (Version 0.2; Müller, 2017), *car* (Version 2.1.4; Fox & Weisberg, 2011), *caTools* (Version 1.17.1; Tuszynski, 2014), *corrplot* (Version 0.84; Wei & Simko, 2017), *cowplot* (Version 0.9.2; Wilke, 2017), *doBy* (Version 4.5.15; Højsgaard & Halekoh, 2016), *dplyr* (Version 0.7.4; Wickham et al., 2017), *emmeans* (Version 1.1.2; Lenth, 2018), *Formula* (Version 1.2.1; Zeileis & Croissant, 2010), *gdata* (Version 2.18.0; Warnes et al., 2017), *ggplot2* (Version 2.2.1; Wickham, 2009), *Hmisc* (Version 4.1.1; Harrell Jr, Charles Dupont, & others., 2018), *knitr* (Version 1.20; Xie, 2015), *lattice* (Version 0.20.35; Sarkar, 2008), *lme4* (Version 1.1.13; Bates, Mächler, Bolker, & Walker, 2015), *MASS* (Version 7.3.47; Venables & Ripley, 2002), *Matrix* (Version 1.2.12; Bates & Maechler, 2017), *multcomp* (Version 1.4.8; Hothorn, Bretz, & Westfall, 2008), *mvtnorm* (Version 1.0.7; Genz & Bretz, 2009), *papaja* (Version 0.1.0.9655; Aust & Barth, 2017), *plyr* (Wickham, 2011; Version 1.8.4; Wickham et al., 2017), *reshape* (Version 0.8.7; Wickham, 2007), *shiny* (Version 1.0.5; Chang, Cheng, Allaire, Xie, & McPherson, 2017), *stringi* (Version 1.1.6; Gagolewski, 2017), *survival* (Version 2.41.3; Terry M. Therneau & Patricia M. Grambsch, 2000), *TH.data* (Version 1.0.8; Hothorn, 2017), *tidyr* (Version 0.6.3; Wickham, 2017), and *xtable* (Version 1.8.2; Dahl, 2016). The data and code used to generate this manuscript (including analyses and plots) can be found at https://osf.io/wm3t6/.

## Results

### Behavioral measures

Mean response times and percent correct are shown as a function of working memory load for each task (working memory and visual search) in figure 4. Paired samples T-tests were carried out to test the effect of working memory load on response times (ms) and accuracy (%) in both the working memory and visual search tasks. The results of the T-tests showed that the high load condition did result in both significantly slower (*M_d_* = 83.25, 95% CI [33.45, 133.05], *t*(19) = 3.50, *p* = .002) and less accurate (*M_d_* = −9.30, 95% CI [−13.83, −4.77], *t*(19) = −4.29, *p* < .001) responses to the working memory task. The results also showed that participants were significantly slower on the visual search task under high working memory load (*M_d_* = 11.10, 95% CI [4.21, 17.99], *t*(19) = 3.37, *p* = .003) however there was no significant effect of working memory load on visual search accuracy (*M_d_* = −0.25, 95% CI [−1.29, 0.79], *t*(19) = −0.50, *p* = .621). These data confirm firstly that the working memory task effectively manipulated the load on a participant’s working memory, and secondly that this manipulation also effected performance on a concurrent visual search task.

**Figure 4.**
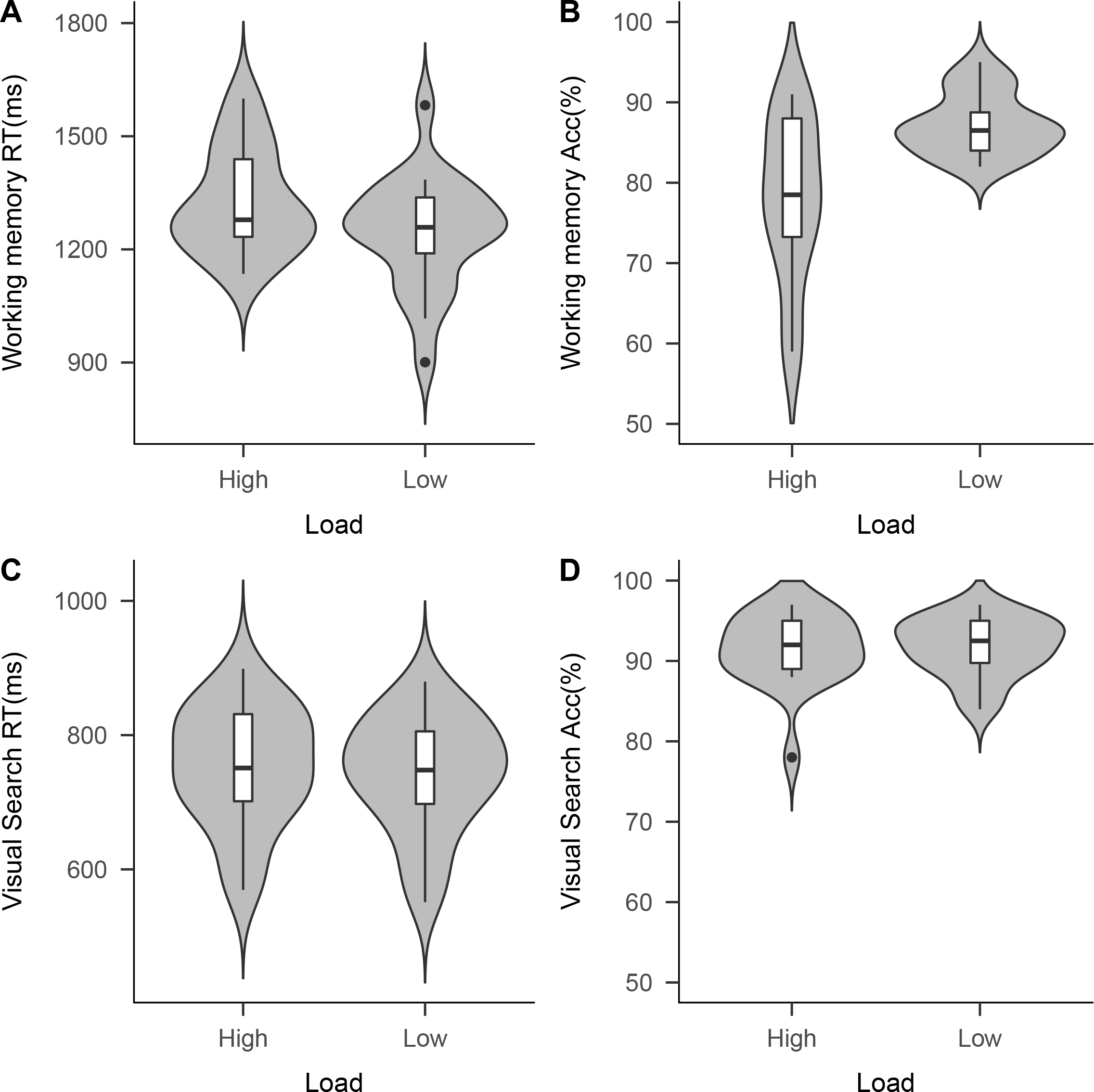
Violin plots showing the effect of working memory load on A) Working memory response times, B) working memory accuracy, C) visual search response times, and D) visual search accuracy.

### Within-subject electrophysiological measures

In order to test for the effect of working memory load on lateralized ERP components related to target and distractor processing, two repeated measures ANOVAs were performed. For lateralized targets, the repeated measures factors were working memory load (high and low), and target component (P1pc, N2pc, SPCNa, and SPCNb) whereas for lateralized distractors the repeated measures factors were working memory load (high and low), and distractor component (N2pc, Ptc, SPCNa, and SPCNb). Greenhouse-Geisser corrected values are used in all cases. The results for targets showed a significant main effect of component (*F*(2.2, 41.81) = 45.22, *MSE* = 0.70, *p* < .001,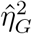 = .409), but no significant main effect of working memory load (*F*(1, 19) = 1.21, *MSE* = 1.02, *p* = .286,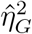 = .012), nor any significant interaction between working memory load and component (*F*(2.57, 48.91) = 2.38, *MSE* = 0.34, *p* = .090,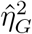 = .020). Pairwise comparisons using Tukey correction were performed in order to follow up the main effect of component. The results showed that the P1pc is significantly more positive in amplitude (M = 0.55, SE = 0.09) when compared to the N2pc (M = −0.71, SE = 0.15, p < 0.001), SPCNa (M = −1.22, SE = 0.19, p < 0.001), and SPCNb (M = −0.77, SE = 0.13, p < 0.001). The N2pc is significantly smaller in amplitude than the SPCNa (p < 0.001), but there is no significant difference in comparison to the SPCNb (p > 0.05). Finally, there is also no significant difference between SPCNa and SPCNb (p > 0.05). Full results shown in table 2.

**Table 1.**
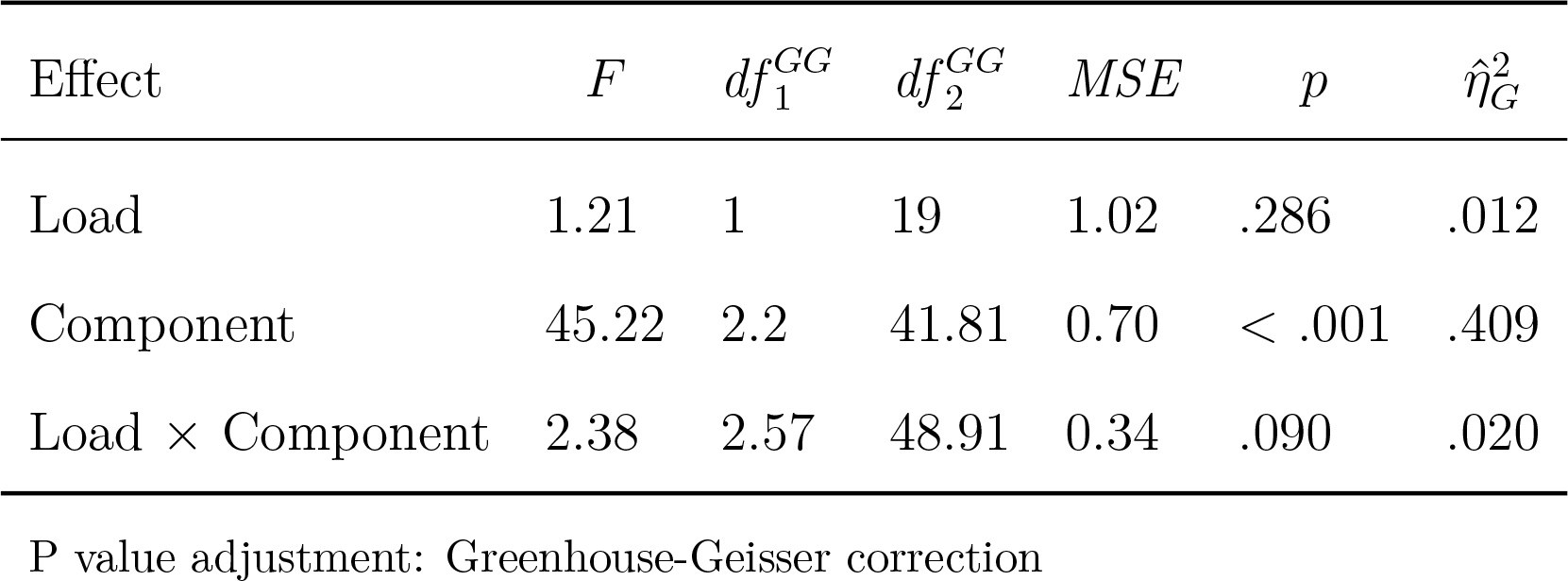
Summary of the 2 × 4 repeated measures ANOVA used to test the effects of working memory load and component on the lateraliesd neural response to targets during visual search.

**Table 2.** Summary of the posthoc comparisons used to follow up the main effect of component on the lateraliesd neural response to targets during visual search.

| contrast | estimate | SE | df | t.ratio | p.value |
| --- | --- | --- | --- | --- | --- |
| P1pc - N2pc | 1.2575 | 0.1600 | 57 | 7.859 | <.0001 |
| P1pc - SPCNa | 1.7732 | 0.1600 | 57 | 11.082 | <.0001 |
| P1pc - SPCNb | 1.3217 | 0.1600 | 57 | 8.261 | <.0001 |
| N2pc - SPCNa | 0.5157 | 0.1600 | 57 | 3.223 | 0.0110 |
| N2pc - SPCNb | 0.0642 | 0.1600 | 57 | 0.402 | 0.9779 |
| SPCNa - SPCNb | -0.4515 | 0.1600 | 57 | -2.822 | 0.0323 |

P value adjustment: tukey method for comparing a family of 4 estimates

The results for distractors showed a significant main effect of component (*F*(1.9, 36.15) = 41.52, *MSE* = 0.74, *p* < .001,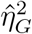 = .481), and a significant main effect of working memory load (*F*(1, 19) = 4.56, *MSE* = 0.69, *p* = .046,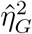 = .048) indicating that the response to distractors under high load is significantly more negative (M=-0.44, SD = 0.97) than the response to distractors under low load (M = −0.15, SD = 0.97). There was no significant interaction between working memory load and component (*F*(1.88, 35.63) = 1.37, *MSE* = 0.29, *p* = .266,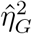 = .012). Pairwise comparisons using Tukey correction were performed in order to follow up the main effect of component. The results showed that the Ptc is significantly more positive in amplitude (M = 0.75, SE = 0.13) when compared to the N2pc (M = −0.63, SE = 0.09, p < 0.001), SPCNa (M = −0.74, SE = 0.13, p < 0.001), and SPCNb (M = −0.56, SE = 0.11, p < 0.001). There were no other significant differences between the N2pc, SPCNa, and SPCNb (see table 4). In order to check for specifity of the effect we performed pairwise comparisons which compared the amplitude of each component under high versus low load. These comparisons suggest that the only component which was significantly affected by working memory load was the Ptc which was smaller under high load (see table 5) however the lack of an interaction between load and component in the ANOVA suggests this specific effect could be accounted for by a generally more positive lateralized response to the distractor under low load.

**Table 3.** Summary of the 2 × 4 repeated measures ANOVA used to test the effects of working memory load and component on the lateraliesd neural response to distractors in the visual search task.

| Effect | $F$ | $df_1^{GG}$ | $df_2^{GG}$ | $MSE$ | $p$ | $\hat{\eta}_G^2$ |
| --- | --- | --- | --- | --- | --- | --- |
| Load | 4.56 | 1 | 19 | 0.69 | .046 | .048 |
| Component | 41.52 | 1.9 | 36.15 | 0.74 | < .001 | .481 |
| Load × Component | 1.37 | 1.88 | 35.63 | 0.29 | .266 | .012 |

P value adjustment: Greenhouse-Geisser correction

**Table 4.** Summary of the posthoc comparisons used to follow up the main effect of component on the lateraliesd neural response to targets during visual search.

| contrast | estimate | SE | df | t.ratio | p.value |
| --- | --- | --- | --- | --- | --- |
| N2pc - Ptc | -1.3735 | 0.1531 | 57 | -8.969 | <.0001 |
| N2pc - SPCNa | 0.1095 | 0.1531 | 57 | 0.715 | 0.8907 |
| N2pc - SPCNb | -0.0657 | 0.1531 | 57 | -0.429 | 0.9732 |
| Ptc - SPCNa | 1.4830 | 0.1531 | 57 | 9.684 | <.0001 |
| Ptc - SPCNb | 1.3078 | 0.1531 | 57 | 8.539 | <.0001 |
| SPCNa - SPCNb | -0.1752 | 0.1531 | 57 | -1.144 | 0.6639 |

P value adjustment: tukey method for comparing a family of 4 estimates

**Table 5.**
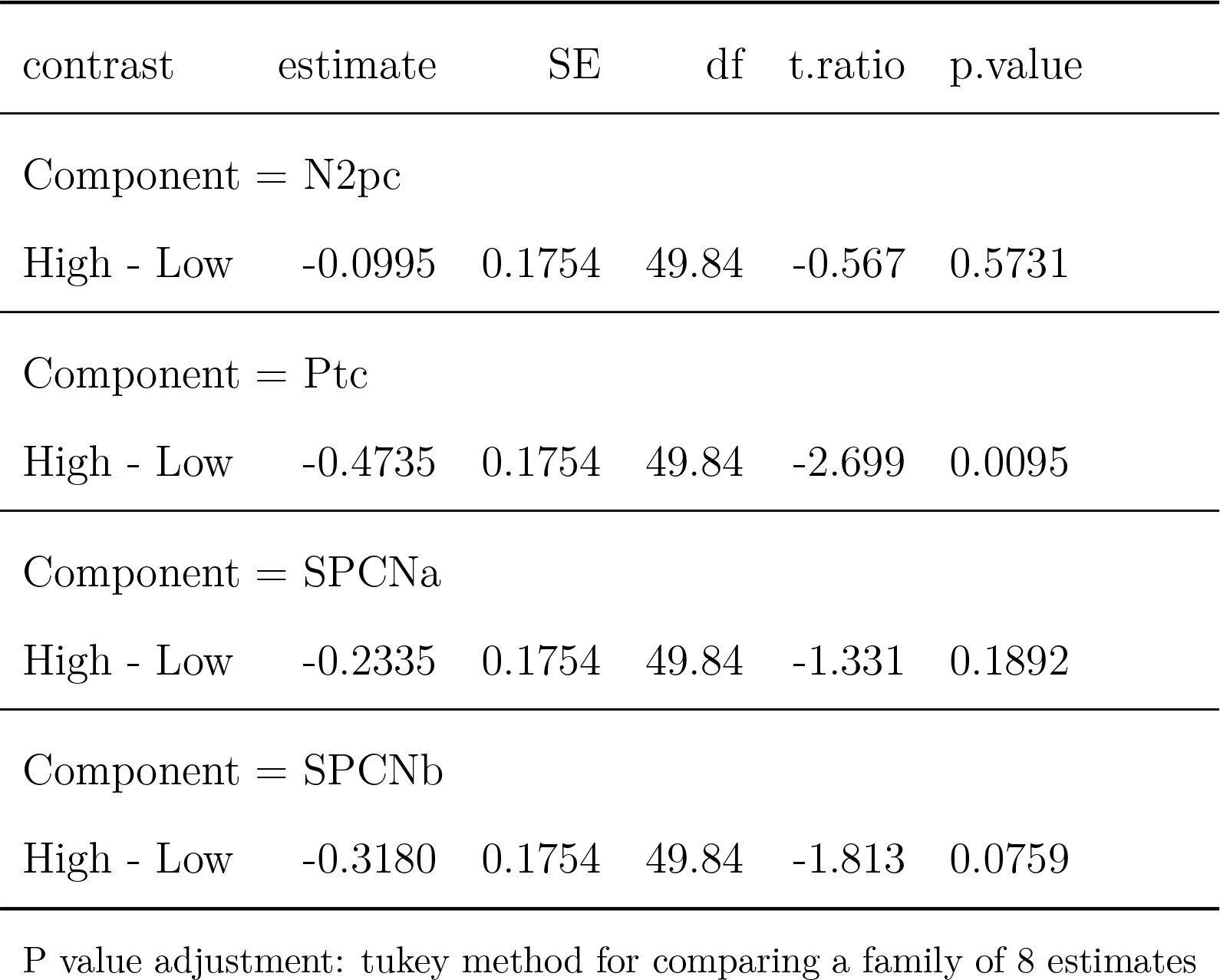
Pairwise comparisons showing the effect of load on each component elicited by lateralized distractors using Tukey correction.

| contrast | estimate | SE | df | t.ratio | p.value |
| --- | --- | --- | --- | --- | --- |
| Component = N2pc |  |  |  |  |  |
| High - Low | -0.0995 | 0.1754 | 49.84 | -0.567 | 0.5731 |
| Component = Ptc |  |  |  |  |  |
| High - Low | -0.4735 | 0.1754 | 49.84 | -2.699 | 0.0095 |
| Component = SPCNa |  |  |  |  |  |
| High - Low | -0.2335 | 0.1754 | 49.84 | -1.331 | 0.1892 |
| Component = SPCNb |  |  |  |  |  |
| High - Low | -0.3180 | 0.1754 | 49.84 | -1.813 | 0.0759 |
| P value adjustment: tukey method for comparing a family of 8 estimates |  |  |  |  |  |

### Between-subject electrophysiological measures

In order to investigate the relationship between lateralized activity and behavior in the tasks, a correlation matrix was made for each working memory load condition. Each matrix contained the behavioral measures (accuracy and response time) on both tasks (working memory task and visual search task), as well as the total lateralized positive area for each stimulus (targets and distractors) and the total lateralized negative area for each stimulus. Under low working memory load, there is a significant correlation between visual search response times and the positivity to targets, positivity to distractors, and the negativity to distractors. Individuals with larger positivities to targets appear to perform faster on the visual search task (r = −0.507, p = 0.023) as do individuals with larger positivities to distractors (r = −0.603, p = 0.005). Individuals with a larger negativity to distractors however, perform slower (r = 0.447, p = 0.048). There is also a significant correlation between working memory response times and the distractor negativity such that individuals with larger negativities to distractors perform slower on the working memory task (r = 0.543, p = 0.013). Scatter plots for these relationships are shown in figure 5. Under high load however, these effects disappear and the only significant relationship between lateralized amplitude and behavior shows that individuals with a larger target negativity tend to perform more accurately on the task (r = 0.538, p = 0.014). While we have only reported on the relationships between amplitudes and behavior here, all significant correlations are visualized for the low load condition in figure 7 and for the high load condition in figure 8. For the full correlation matrix for the low load condition see table 6, and for the high load condition see table 7.

**Figure 5.**
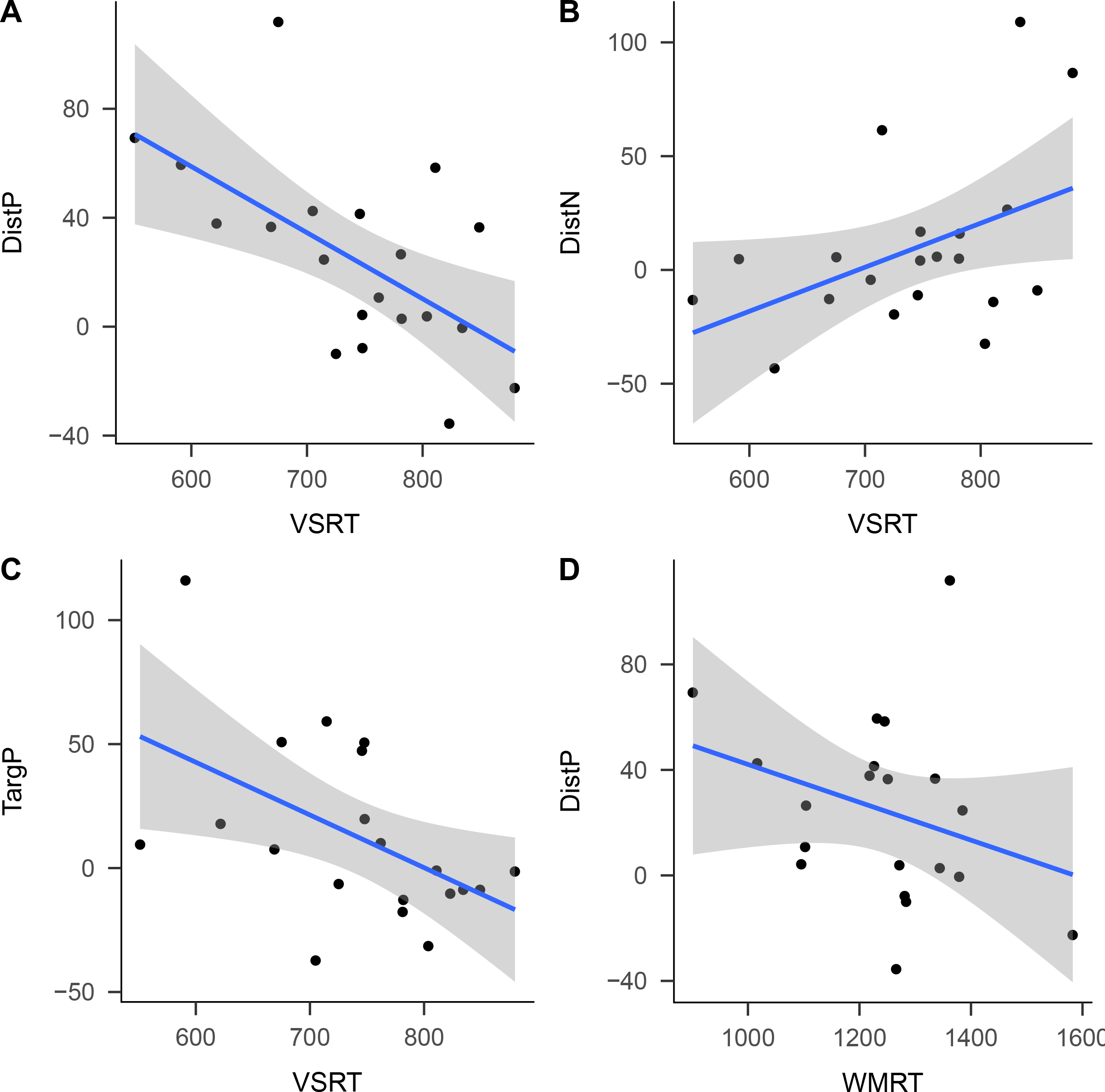
Scatter plots showing the significant relationships between neural responses and behavior in the low working memory load condition. A) visual search response times and distractor positivity, B) visual search response times and distractor negativity, C) visual search response times and target positivity, and D) working memory response times and distractor positivity.

**Figure 6.**
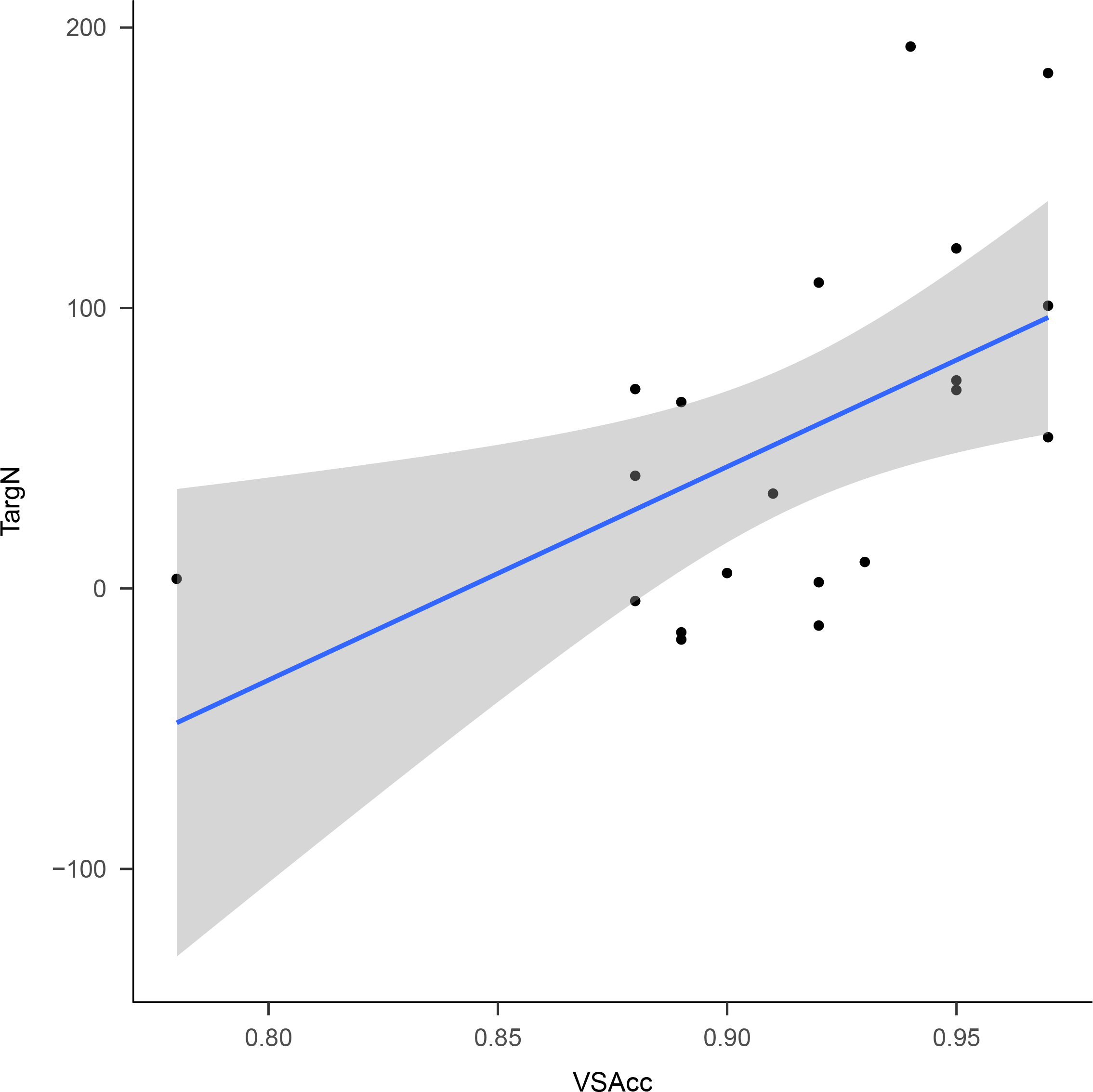
Scatter plot showing the significant relationship between visual search accuracy and target negativity in the high working memory load condition.

**Figure 7.**
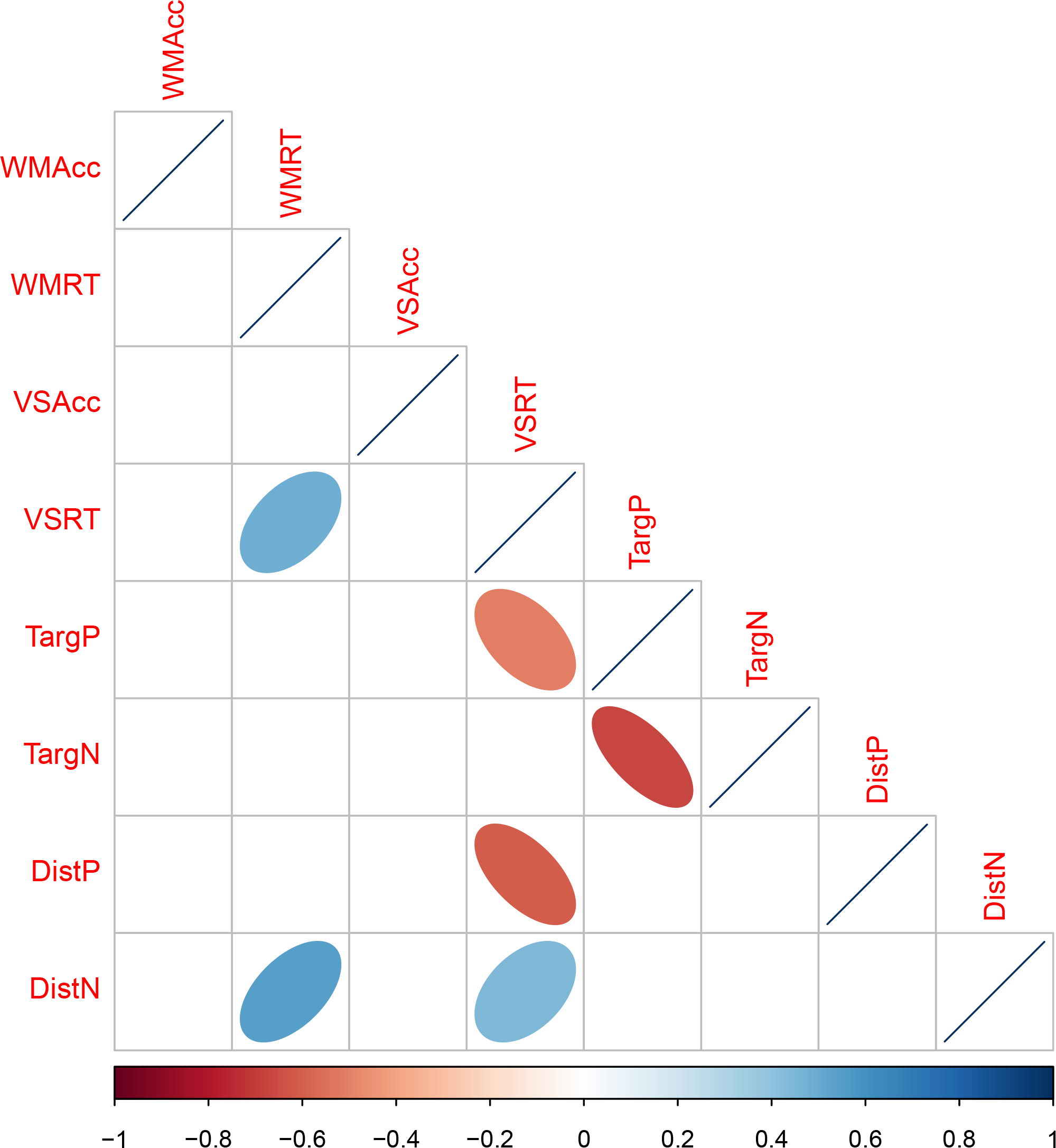
Correlogram showing the direction and strength of all significant correlations between behavioral and neural measures under low working memory load. WM = working memory task, VS = visual search task, Acc = accuracy, RT = response time, Targ = lateralized target response, Dist = lateralized distractor response, P = positive area, N = negative area.

**Figure 8.**
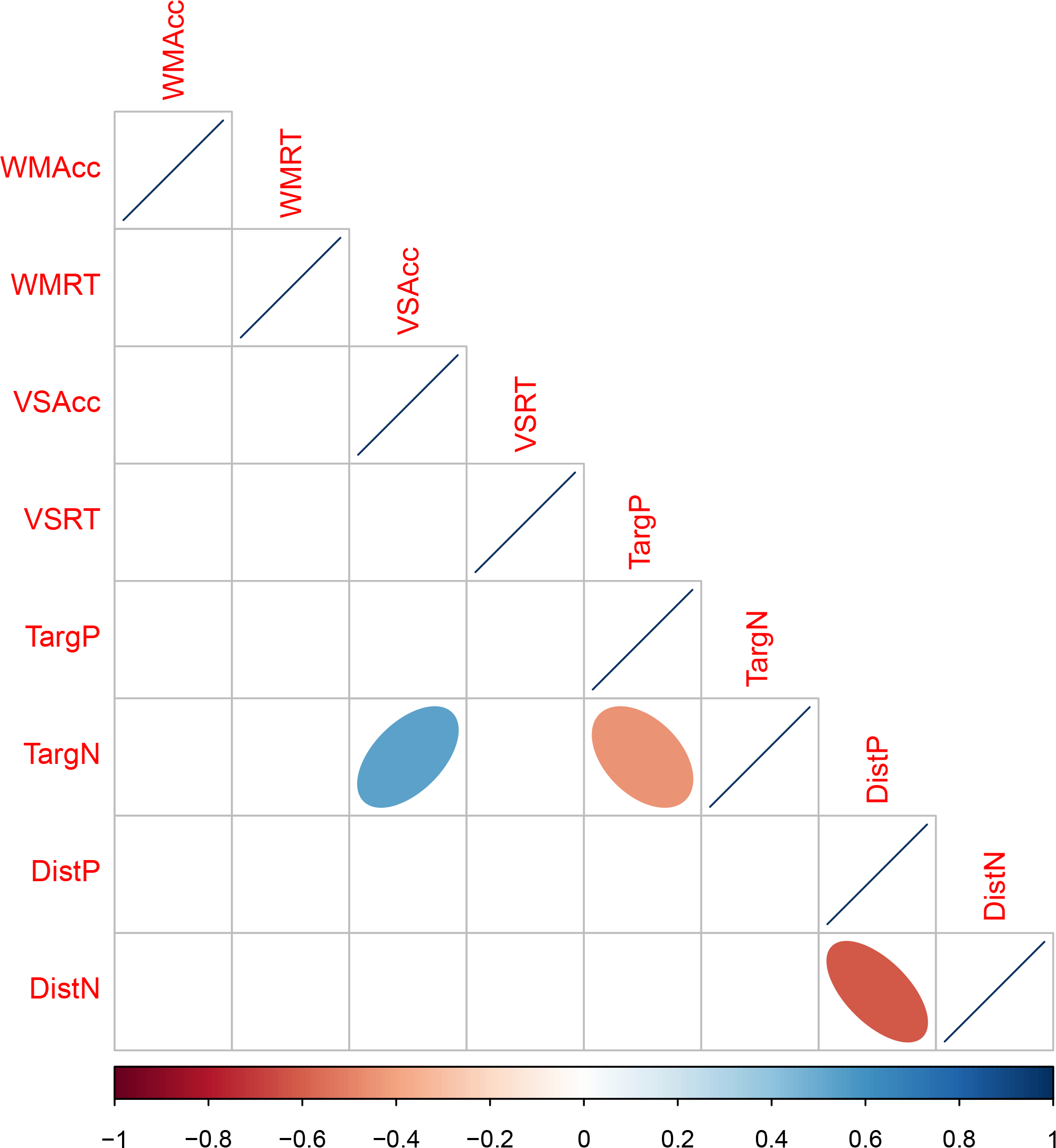
Correlogram showing the direction and strength of all significant correlations between behavioral and neural measures under high working memory load. WM = working memory task, VS = visual search task, Acc = accuracy, RT = response time, Targ = lateralized target response, Dist = lateralized distractor response, P = positive area, N = negative area.

**Table 6.** Correlation matrix for the low load condition showing the relationships between all behavioral and ERP measures. WM = working memory task, VS = visual search task, Acc = accuracy, RT = response time, Targ = lateralized target response, Dist = lateralized distractor response, P = positive area, N = negative area.

|  | WMAcc | WMRT | VSAcc | VSRT | TargP | TargN | DistP |
| --- | --- | --- | --- | --- | --- | --- | --- |
| WMAcc |  |  |  |  |  |  |  |
| WMRT | -0.38* |  |  |  |  |  |  |
| VSAcc | 0.16 | 0.07 |  |  |  |  |  |
| VSRT | -0.14 | 0.48** | 0.18 |  |  |  |  |
| TargP | -0.12 | 0.07 | -0.15 | -0.51** |  |  |  |
| TargN | -0.02 | -0.20 | 0.18 | 0.42* | -0.67*** |  |  |
| DistP | 0.23 | -0.31 | -0.06 | -0.60*** | 0.39* | -0.15 |  |
| DistN | -0.14 | 0.54** | 0.40* | 0.45** | 0.04 | -0.07 | -0.38* |

Note: $p < .001$ '\*\*\*\*'; $p < .01$ '\*\*\*'; $p < .05$ '\*\*'; $p < .1$ '\*'

**Table 7.** Correlation matrix for the high load condition showing the relationships between all behavioral and ERP measures. WM = working memory task, VS = visual search task, Acc = accuracy, RT = response time, Targ = lateralized target response, Dist = lateralized distractor response, P = positive area, N = negative area.

|  | WMAcc | WMRT | VSAcc | VSRT | TargP | TargN | DistP |
| --- | --- | --- | --- | --- | --- | --- | --- |
| WMAcc |  |  |  |  |  |  |  |
| WMRT | 0.23 |  |  |  |  |  |  |
| VSAcc | 0.23 | 0.23 |  |  |  |  |  |
| VSRT | -0.10 | 0.37 | 0.35 |  |  |  |  |
| TargP | 0.11 | -0.28 | -0.35 | -0.43* |  |  |  |
| TargN | 0.12 | 0.05 | 0.54** | -0.11 | -0.45** |  |  |
| DistP | 0.20 | 0.00 | 0.38* | 0.20 | -0.02 | 0.12 |  |
| DistN | -0.24 | 0.29 | -0.25 | 0.10 | -0.02 | -0.17 | -0.62*** |

Note: $p < .001$ '\*\*\*\*'; $p < .01$ '\*\*\*'; $p < .05$ '\*\*'; $p < .1$ '\*'

## Discussion

The aim of this research was to specify the relationship between selective attention and working memory using neural indices of attentional control. Our results showed that within individuals, increased working memory load slowed responses on a concurrent visual search task. The lateralized neural response to targets and distractors in the visual search displays showed that this appeared to be the result of changes in the way salient distractors were processed as working memory load increased. Target processing on the other hand appeared unaffected by working memory load. The presence of a main effect of load on distractors, but no interaction between load and component means that it is difficult to specify the attentional process (whether capture by distractors, or disengagement from distractors) being impacted. Follow-up analyses suggest that the best candidate is the distractor Ptc which indexes distractor disengagement and is significantly reduced under high working memory load. Taken together, these results support the idea that working memory load specifically impacts an individual’s ability to disengage from salient distractors, and that failure to efficiently disengage from distractors is responsible for the behavioral impairments observed under increasing working memory load (Fukuda & Vogel, 2011).

The effect of working memory load on the neural response to distractors raises some interesting questions about the dynamics of lateralized positivities recorded in this task and others. Our results showed that as working memory load was increased, the neural response to distractors became less positive across all four of the components we measured. If each of these components reflects an independent stage of processing (Gaspar & McDonald, 2014; Hickey et al., 2009; Hilimire et al., 2011), then one explanation for our result is that increased working memory load impacted on distractor capture, distractor disengagement, and storage of distractor features in such a way that the neural index of these processes became less positive. When the components were tested individually however, our data showed that only the Ptc was significantly modulated by working memory load. A parsimonious explanation for our results is that increased memory load impacted on the amplitude of the distractor Ptc, but that Ptc activity overlapped in time with the other components of the average ERP. Thus, any modulation of Ptc amplitude would be observed not only at the peak of the Ptc itself, but in the amplitude of all overlapping activity. The apparent effect of overlap that we observe in our results raises the question of why overlap between components would be observed in our experiment when previous research has shown these components to be dissociable.

The reason for differences in the extent to which Ptc overlaps with other components depends on how Ptc manifests at the single trial level. A sustained positivity in the average ERP could reflect either a sustained positive component at the single trial level, or variability in the latency of the component at the single trial level. In the former case, Ptc reflects the presence of a sustained lateralized positivity, the bulk of which is typically masked by a series of overlapping, punctuated, lateralized negativities. Differences across experiments in the extent of observable overlap could therefore be the result of differences in the proportion of negative to positive activity in the task. Alternatively, the lateralized positivity we recorded may be short in duration but highly variable in latency across trials, therefore the sustained positivity in the average waveform is the product of convolving this variable signal across the epoch. Differences in overlap across experiments in this case could instead be due to differences in the amount of latency variability induced by the specific task. In both cases, the interpretation of our results remains the same; that increased working memory load impacts specifically on the amplitude of the lateralized positivity to distractors, and that this reflects impairments of distractor disengagement. Our experiment didn’t allow for teasing apart these two scenarios, nonetheless our result does support the use of signed area approaches for quantifying the amplitude of these lateralized positivities as suggested by Sawaki et al. (2012). Whether the lateralized positivity is a large sustained component, or highly variable across trials, the signed-area approach would be well suited to capturing and quantifying its amplitude.

With respect to individual differences in performance on the visual search task and the working memory task, the effects are broader. Under low load, individuals who perform faster on the visual search task appeared to have larger lateralized positivities in response to both targets and distractors, as well as smaller lateralized negativities to distractors. This finding indicates that stimulus disengagement is an important individual characteristic related to response times under load. Whether an object is a target or distractor, the larger the disengagement signal, the faster participants responded in the visual search task. It’s worth noting that faster response times do not appear to come at the expense of accuracy on the task as demonstrated by a lack of correlation between speed and accuracy. The correlation with distractor negativities also supports the notion that individuals who attend to and store the distractor representation less, gain an advantage in target response times. Under high load, the relationships between distractor processing and performance as well as target positivity and performance disappear. Instead we observe that increased target negativity is related to more accurate responses on the visual search task. These individual differences are consistent with the within-subject effects described above which show that increased lateralized positivities are associated with better performance. The between-subject findings however introduce the idea that target disengagement is an important factor for distinguishing individuals.

Individual differences in working memory capacity and its relationship to lateralized ERP components has also been carried out previously by Gaspar et al. (2016). In their experiment, working memory capacity was measured in a change detection task that was performed separate to the visual search task used to evoke the lateralized components. Gaspar et al. (2016) showed that positivity to distractors was correlated with working memory capacity yet there was no evidence for a relationship between target processing and working memory capacity. At first glance, our between subjects analyses appear to contradict the results of Gaspar et al. (2016). We found relationships between individual performance and the processing of both targets and distractors. However, differences in both task design and analysis between our study and that of Gaspar et al. (2016) could account for differences in the results. For instance, our study explored correlations between component amplitudes and performance on the visual search task, not the working memory task. In fact, when we looked at the relationship between component amplitude and working memory performance, the only significant effect we observed was between distractor processing and working memory response times which mirrors the results of Gaspar et al. (2016). Gaspar and colleagues also correlated ERP measures and performance across two independent tasks, whereas the tasks in our study were performed concurrently. Given the changes we observed in our correlations across load conditions, we wouldn’t expect our correlational findings to be consistent with those from the task used by Gaspar et al. (2016) (which would represent a no-load condition).

Unexpectedly we detected an early positivity in the lateralized response to targets. Although the P1pc has been observed in other paradigms, its functional role is less well understood. Current hypotheses to explain the occurrence of a P1pc don’t seem to apply to our findings. One feature of visual search tasks that can elicit a P1pc is sensory imbalance across the visual display (Edwards & Drasdo, 1987). If a luminant stimulus is presented in one visual field without a luminance matched stimulus in the opposite visual field location then we would observe an increased P1 contralateral to the lateralized target. This would manifest in the lateralized ERPs as P1pc. In our task, all targets were presented opposite gray fillers which were physically matched for luminance. Although this is not guaranteed to prevent the target color from eliciting a larger P1, the lack of any P1pc for distractors in our task (which had the exact same colors as targets and in the same proportions) suggests that the P1pc is not a result of stimulus imbalance. Alternatively, growing evidence suggests that P1pc may be elicited in response to stimulus change (Verleger, Grajewska, & Jaskowski, 2012). Although the target color changed across trials in our study, this is unlikely to explain the P1pc in our task. First, because target color changed at the exact same frequency as distractor color which did not elicit a P1pc. Second, the target-decoy task used in our study is a replication of the task used by Hilimire et al. (2011) who did not record a P1pc to targets. The P1pc we observe in our task is therefore more likely to be a result of the concurrent working memory task.

One explanation for how the working memory task could elicit a target P1pc in our target-decoy task is that the P1pc that we measured in our task is functionally distinct from the P1pc that has been investigated by other researchers (Verleger et al., 2012). The P1pc in our task occurred slightly later than the typically observed P1pc in other studies and may be related to the process of stimulus disengagement that we associate with the Ptc. Support for this explanation comes from similarities between the target positivity and distractor positivity in terms of their relationships to other measures in the task. For example, target positivity was inversely correlated with target negativity. This was also the case for distractor positivity and distractor negativity. Target positivity was also inversely correlated with visual search response times under low load but not under high load, mirroring the correlations we saw between distractor positivity and behavior. Another source of evidence for our explanation that target positivity in our task reflects disengagement is that under low load, the correlation between target and distractor positivity is approaching significance (see table 6). This finding indicates that individuals with a larger distractor positivity also tend to have a larger target positivity. Although previous studies have found a Ptc elicited by targets, it has not been observed in as early a time window as in our study. The role of a concurrent working memory task in eliciting an early, target related positivity, and the functional significance of this component requires further investigation.

## Conclusion

This research attempted to answer the question of why working memory availability is important for selective attention. In summary, the results suggest that working memory availability enables individuals to disengage from salient distractors which have captured attention. Our findings are consistent with evidence from between-subject designs showing that individuals with low working memory capacity have specific impairments in distractor processing (Fukuda & Vogel, 2011; Gaspar et al., 2016). Our results extend this work to show that when working memory load is increased on an individual (and therefore their working memory availability is reduced), it is similarly their ability to disengage from salient distractors which is impaired. Our results also raise questions for future research about the extent and effect of overlap between lateralized ERP components, as well as the cause and the significance of an early lateralized positivity to targets. Taken together, the results suggest that a fruitful approach for characterizing the relationship between working memory and selective attention is to focus on the availability of working memory resources to unify between-subject and within-subject comparisons. We also demonstrate the utility of lateralized event-related potentials in teasing apart sub-processes of selective attention which are diffcult to dissociate behaviorally.

